# Mate choice confers direct benefits to females of *Anastrepha fraterculus* (Diptera: Tephritidae)

**DOI:** 10.1101/584128

**Authors:** Guillermo E. Bachmann, Francisco Devescovi, Ana L. Nussenbaum, Fabián H. Milla, Todd E. Shelly, Jorge L. Cladera, Patricia C. Fernández, María T. Vera, Diego F. Segura

## Abstract

Exposure to plant compounds and analogues of juvenile hormone (JH) increase male mating success in several species of tephritid fruit flies. Most of these species exhibit a lek mating system, characterized by active female choice. Although the pattern of enhanced male mating success is evident, few studies have investigated what benefits, if any, females gain via choice of exposed males in the lek mating system. In the South American fruit fly, *Anastrepha fraterculus*, females mate preferentially with males that were exposed to volatiles released by guava fruit or treated with methoprene (a JH analogue). Here, we tested the hypothesis that female choice confers direct fitness benefits in terms of fecundity and fertility. We first carried out mate choice experiments presenting females with males treated and non-treated with guava volatiles or, alternatively, treated and non-treated with methoprene. After we confirm female preference for treated males, we compared the fecundity and fertility between females mated with treated males and non-treated ones. We found that *A. fraterculus* females that mated with males exposed to guava volatiles showed higher fecundity than females mated to non-exposed males. On the other hand, females that mated methoprene-treated males showed no evidence of direct benefits. Our findings represent the first evidence of a direct benefit associated to female preference for males that were exposed to host fruit odors in tephritid fruit flies. Differences between the two treatments are discussed in evolutionary and pest management terms.

## Introduction

Tephritid fruit flies (Diptera: Tephritidae) infest hundreds of different plant species, including many economically important fruits [1]. One environmentally friendly control strategy used against fruit fly pests is the sterile insect technique (SIT) [2], which requires solid knowledge of the sexual behaviour of the target pest [3]. SIT is based on the ability of mass-released, sterile males to mate with fertile, wild females and hence induce sterility in the pest population [4]. Many species of tephritid fruit flies exhibit lek mating systems characterized by female choice, and this has prompted considerable attention on the factors that influence male mating success [5,6]. The collective outcome of this research has been the identification of several pre-release treatments that boost the sexual competitiveness of male tephritids [7–10] including: pre-release diet containing a proteinaceous source [11–20]; exposure of males to semiochemicals [9,10,21,22]; and the use of methoprene, a juvenile hormone (JH) analogue, to both boost male mating success and accelerate their sexual maturation [7,17,18,23].

Although the drive to improve SIT has, in many cases, identified male traits associated with mating success, little is known about the ecological and evolutionary forces that shape female mate preferences, particularly the potential fitness benefits associated with mate choice [24]. Kirkpatrick and Ryan [25] and Wyatt [26] proposed that female choice is generally associated with direct benefits, such as increased fecundity and longevity, reduced risk of predation, or access to material resources under male control [27–29]. Simultaneously, females may gain indirect benefits if mate choice positively affects the fitness of their offspring, such as those explained under the sexy son and good genes hypotheses [30–33]. For tephritids, female choice has been associated with direct and indirect benefits [34,35]. In particular, the association between female benefits and mate selection mediated via feeding or exposure of males to plant-borne compounds has been investigated in four tephritid species: *Bactrocera dorsalis* (Hendel) [36,37]; *Bactrocera tryoni* (Froggatt) [38,39]; *Ceratitis capitata* (Wiedemann) [13,40]; and *Zeugodacus cucurbitae* (Coquillett) [24]. Although females of this species showed a preference for males either exposed or fed with plant-derived compounds, only Kumaran *et al*. [38,39] found evidence of direct and indirect benefits, respectively, for *B. tryoni* females.

The South American fruit fly, *Anastrepha fraterculus* (Wiedemann), is a polyphagous species that attacks more than 100 species of fruit plants [41], many of which have a high commercial value. Vera *et al*. [42] found an increase in the mating success of *A. fraterculus* males following exposure to the volatiles of guava fruits (*Psidium guajava* L.), a native plant and one of their main hosts in the wild. This phenomenon was later confirmed and extended by Bachmann *et al*. [43], who also found that males exposed to guava fruit volatiles released larger amounts of sex pheromone and performed courtship behaviours more frequently than non-exposed males. Similarly, topical applications of methoprene conferred a mating advantage to the males which also seems to be associated to larger amounts of pheromone being released by methoprene-treated males when competing with non-treated males [44].

Despite evidence showing that guava fruit volatiles and mimics of natural compounds (like methoprene) enhance male mating success in different *Anastrepha* species, no prior investigation has measured the potential fitness benefits gained by females that mate with those enhanced males. Even though guava volatiles are natural compounds and methoprene is synthetic, they both stimulate males to release larger amounts of pheromone and increased male mating success [43,44]. Furthemore, plants produce several natural analogues of the juvenile hormone (termed juvenoids), which protect them from herbivore larvae [45–47]. According to Bede & Tobe [48], some species of insects have adapted to consume these juvenoids to increase their own reproductive output. So, even though methoprene is a synthetic compound, it could still trigger natural responses in males (enhanced signalling) and females (attraction to such signalling). In the present work, we evaluated whether the preference of *A. fraterculus* females for males exposed to guava volatiles and those treated with methoprene derive from direct benefits in terms of increased fecundity and fertility. Because in *A. fraterculus* longer copulations result in longer refractory periods [49], which can decrease the need for future mates and consequently the costs in terms of energy and re-exposure to predators [50], we also evaluated copula duration as potential direct benefit to females.

## Materials and Methods

### Biological material

*Anastrepha fraterculus* flies were obtained from a laboratory colony kept at Instituto de Genética E. A. Favret (IGEAF) which was originally established at Estación Experimental Agroindustrial Obispo Colombres (Tucumán, Argentina) in 1997 with pupae obtained from infested guavas collected in Tafí Viejo (26°43’25’’S 65°16’43’’W, Tucumán, Argentina) [51]. Rearing followed standard procedures using an artificial diet based on yeast, wheat germ, sugar, and agar for larvae [52] and a mixture of sugar, hydrolysed yeast (MP Biomedicals, San Francisco, CA, USA), hydrolysed corn (ARCOR, Tucumán, Argentina) (4:1:1 ratio) and vitamin E (Parafarm, Buenos Aires, Argentina) for adults [53]. Flies used in the tests were virgin and sexually mature (males were > 10 days-old; females were > 14 days-old) [54] and were kept under controlled environmental conditions (24 ± 2 °C, 70 ± 10% RH, and a 12L: 12D photoperiod).

One day after adult emergence flies were sorted by sex, transferred to plastic containers, and provided food and water. Females were placed in 1 L plastic cylindrical containers (15 cm tall, 12 cm in diameter) in groups of 25 individuals and fed with the standard diet. Males were placed in 21 L plastic containers (37 × 28 × 21 cm) in groups of 100 individuals and fed with sugar and brewer’s yeast (CALSA, Tucumán, Argentina) (3:1 ratio).

### Experiments

#### Experiment 1. Mating and reproductive output of females offered males exposed or not exposed to guava volatiles

In order to determine whether the preference of *A. fraterculus* females for guava-exposed males [44] is associated with direct fitness benefits in terms of mating and reproductive output, females were first given the choice to mate with guava-exposed or non-exposed males. Then, we determined differences in mating and reproductive parameters between females that selected exposed or non-exposed males.

Treated males were exposed to guava volatiles without physical access to the fruit following procedures of Bachmann *et al*. [43]. Non-exposed males were kept under the same environmental conditions but in a different room and had no exposure to guava odours. After exposure, males were kept in the 21 L plastic containers in separate rooms and under the same conditions described above. In the mating test, a total of 401 experimental arenas were established, each consisting of three virgin flies: one exposed male, one non-exposed male of the same age, and one female. Males were 12-14 d old; ; whereas females were 14-18 d old. These experimental arenas have been extensively used in *A. fraterculus* as a valid experimental approach to study female mate choice [42–44,55,56]. Males were marked on their thorax with a dot of non-toxic, water-based paint for identification [54]. Randomly assigned colours identified different male treatments. Males and females were released in the experimental arenas early in the morning under semi-darkness. Males were released 15 min before females. Once all arenas were set up, fluorescent lights were turned on, and the occurrence of mating pairs was monitored continuously for 2 h during the natural period of mating activity for Argentinean populations of *A. fraterculus* (9 – 11 am) [54,57]. Whenever a couple was detected, male colour and time at which copulation started and ended were recorded. The recording of copula start and end times was done continuously, and each mating couple was checked every 2-3 minutes. Due to inadvertent errors in recording times, 4 replicates were excluded from the analysis of mating duration. Experiments were conducted under laboratory conditions (25 ± 1 °C and 70 ± 20% RH). Illumination was provided by fluorescent tubes and natural light coming from a window.

After the mating test, mated females were transferred to 3 L glass flasks (with water and food) in groups of three according to the type of male mated. We used groups of females instead of solitary individuals because several studies indicate that the presence of conspecific females stimulates the oviposition [58–60]. Each flask with three females was considered a replicate, with 25 and 22 replicates for females mated to exposed or non-exposed males, respectively. Every 3 days each replicate was provided an oviposition substrate (hereafter, oviposition unit) that consisted of a cylindrical plastic vial (2 cm tall, 2.5 cm in diameter) filled with water coloured with edible red dye (Fleibor, Tablada, Buenos Aires, Argentina) and covered with Parafilm M (Pechiney Plastic Packaging, Chicago, Illinois, USA). This oviposition unit is normally used in the laboratory rearing of *A. fraterculus* at IGEAF. After 24 h, the oviposition unit was removed, and the eggs were recovered using a plastic pipette and then transferred onto a black filter paper. This was, in turn, placed inside a plastic Petri dish (2 cm tall, 8 cm in diameter) on top of a wet cloth and then covered with its lid. The number of eggs from each oviposition unit was counted under a stereoscopic microscope (20x). Petri dishes were then placed inside an incubator at 25 ± 1 °C and 70 ± 10% RH for 4 days to allow embryonic development. After this time, the numbers of hatched and unhatched eggs were counted under a stereoscopic microscope (20x). This entire process was repeated nine times with each replicateduring 27 days, thus covering the peak period of egg-laying in this species [61]. Whenever a dead female was detected, it was removed from the flask and recorded, thus allowing computation of ovipositions *per capita* for each day of egg collection. Due to logistic problems with the incubator, 7 and 1 replicates coming from females mated with exposed and non-exposed males, respectively, were discarded and not considered in the data analysis.

#### Experiment 2. Mating and reproductive output of females offered males treated or not treated with methoprene

To determine whether the preference of *A. fraterculus* females for methoprene-treated males [44] is associated with a direct benefit, we followed a protocol similar to that described for Experiment 1.

The procedure for methoprene application followed Teal *et al*. [62]. Briefly, on the day of emergence males were topically treated by applying 1 µL of a solution of methoprene dissolved in acetone (5 μg/μl) into their torax. Females, treated males, and non-treated males were kept in separate rooms under the same environmental and feeding conditions as described above. In the mating tests a total of 166 experimental arenas were set up (4 replicates were excluded from the analysis of mating duration due to mistakes during the recording of duration times). At the day of the mating test, males were 12 d old and females were 14-18 d old. Afterwards, mated females were transferred in groups of three to cylindrical, 1 L plastic containers. Fecundity and fertility were assessed following the same procedures as described for experiment 1, except that oviposition units were offered seven times to each replicate, covering a total of 21 days. The number of replicates was 24 for females mated to methoprene-treated males and 16 for females mated to non-treated males.

### Data analysis

The numbers of copulations achieved by guava-exposed and non-exposed males (experiment 1) or methoprene-treated and non-treated males (experiment 2) were compared by means of a G test of goodness of fit to an equal proportion hypothesis. The latency to mate (i.e., time elapsed between female release and mating), the mating duration (i.e., time elapsed between the start and end of mating), the overall fecundity (i.e., number of eggs laid across all the egg collections), and the fertility (i.e., average of hatch rate across all egg collections) were compared between females mated to treated or non-treated males by means of a t-test for independent samples. Assumptions of normality of the residuals and homoscedasticity were checked prior to each test. In order to meet the homoscedasticity assumption, mating duration from experiment 1 and fertility in experiment 2 were ln- and logit-transformed, respectively. Also, fertility was logit-transformed in experiment 2. Finally, we carried out a series of survival analyses (Kaplan-Meier estimation, Log-rank test) to evaluate the temporal pattern of female fecundity after mating with exposed or non-exposed males (experiment 1) or treated or non-treated males (experiment 2). STATISTICA 7 [63] and GraphPad Prism 6 [64] were used for statistical analyses and preparation of figures.

## Results

### Experiment 1. Mating and reproductive output of females offered males exposed or not exposed to guava volatiles

A total of 401 mating arenas were established of which 387 resulted in mating. Females mated significantly more often with males exposed to guava than with non-exposed males (G = 19.73, N = 387, d.f. = 1, p < 0.001) (Fig 1). The latency to mate was not affected by the type of male chosen for mating (t = 0.284, d.f. = 385, p = 0.776) (Fig 2a). Mating duration, on the other hand, was longer in matings that involved males exposed to guava volatiles (t = 2.626, d.f. = 381, p < 0.01) (Fig 2b). Female fecundity was significantly different between females mated with exposed and non-exposed males (t = 2.145, d.f. = 45, p = 0.037), as females mated with guava exposed males laid significantly more eggs (Fig 2c). Fertility was not statistically different between the two types of females (t = 0.372, d.f. = 37, p = 0.712) (Fig 2d).

**Fig 1.**
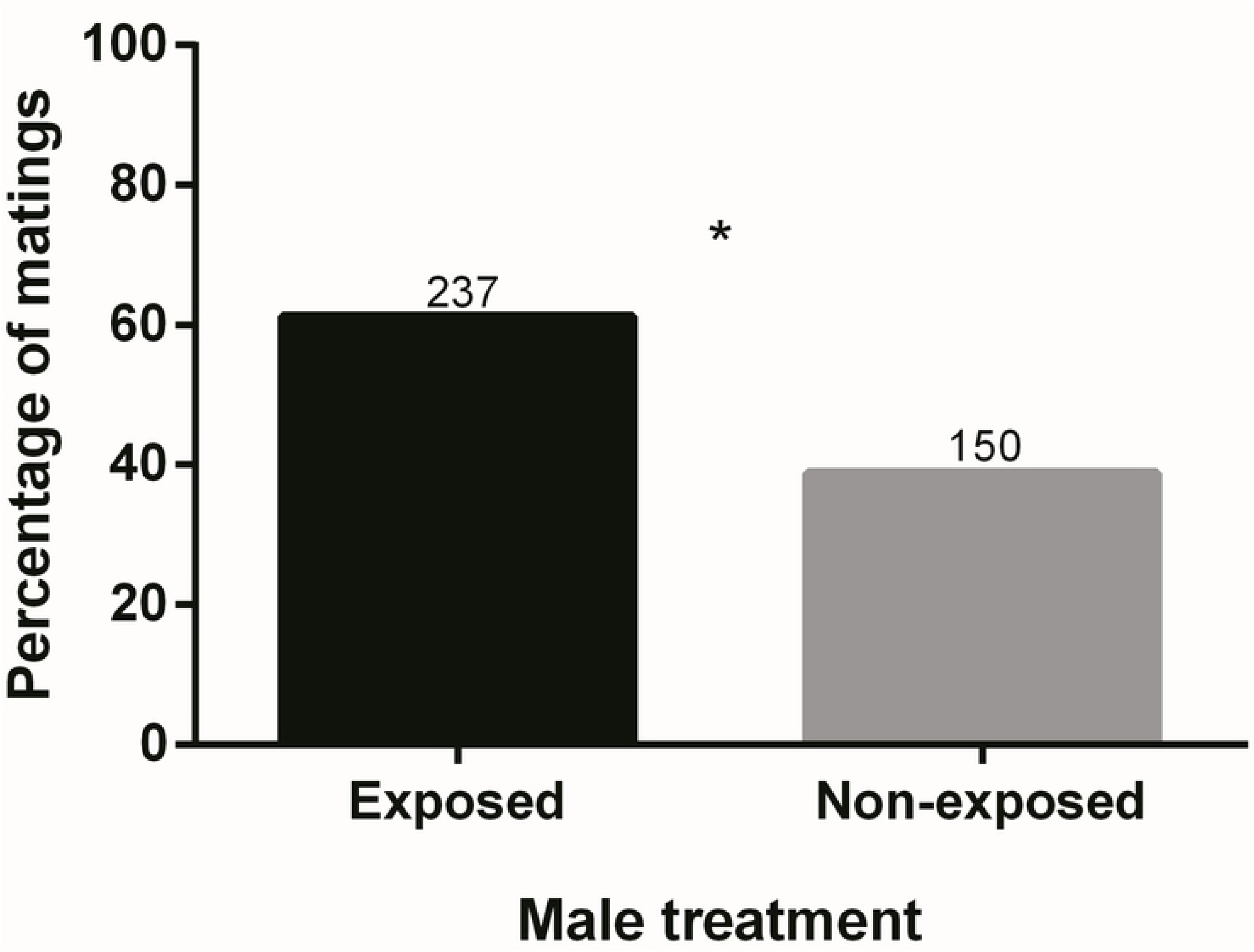
Percentage of matings obtained by males exposed or non-exposed to guava volatiles. Numbers above bars represent numbers of matings. * statistically significant difference (α = 0.05).

**Fig 2.**
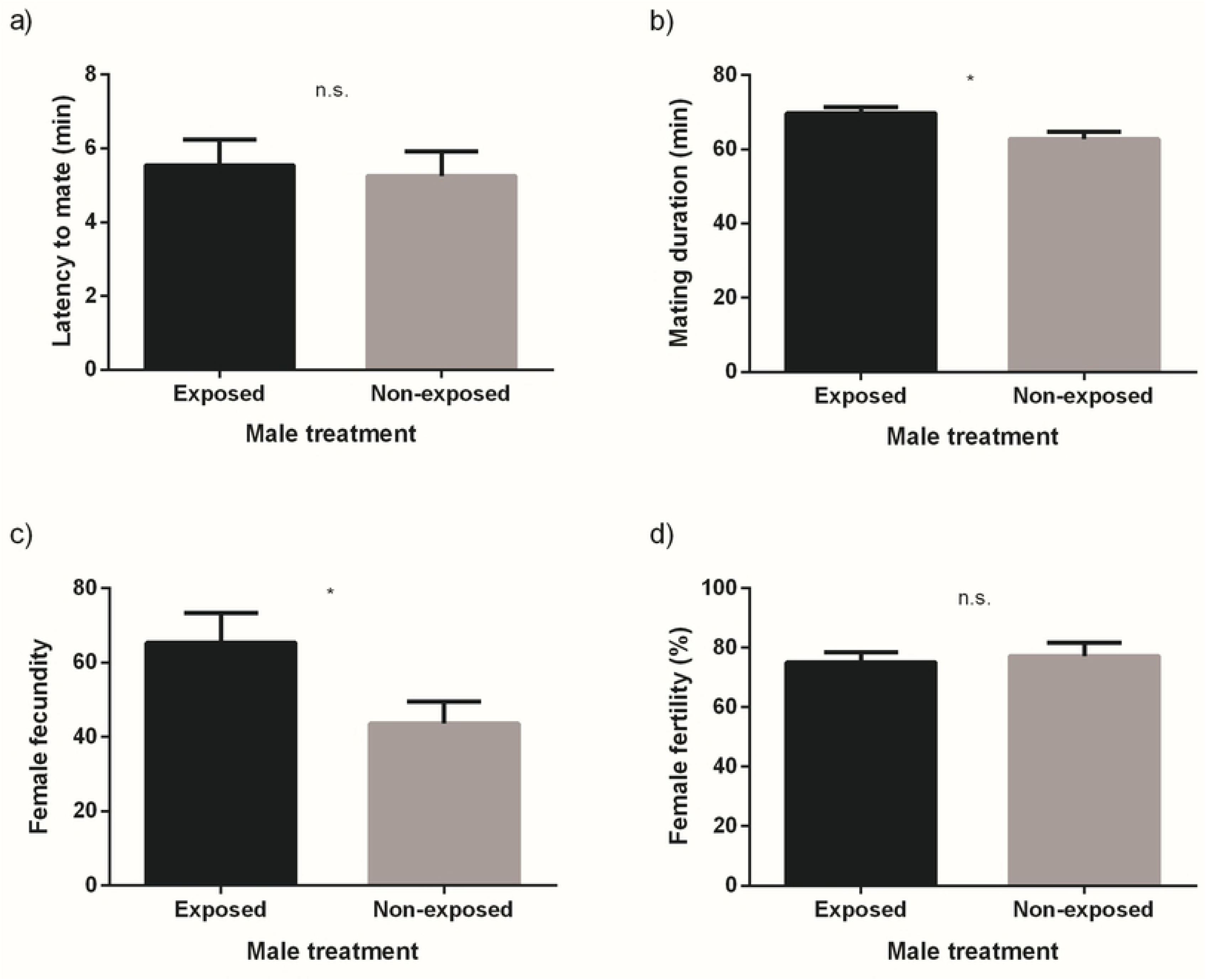
Latency to mate (a), mating duration (b), fecundity (*per capita*) (c) and fertility (d) for females mated to males exposed or non-exposed to guava volatiles (mean and SE). * statistically significant difference (α = 0.05); n.s. not statistically significant difference.

The temporal pattern of egg-laying was independent of the type of male chosen for mating (50% of the total eggs deposited by the 7^th^ irrespective of male type) (χ^2^ = 1.02, d.f. = 1, p = 0.312) (Fig 3).

**Fig 3.**
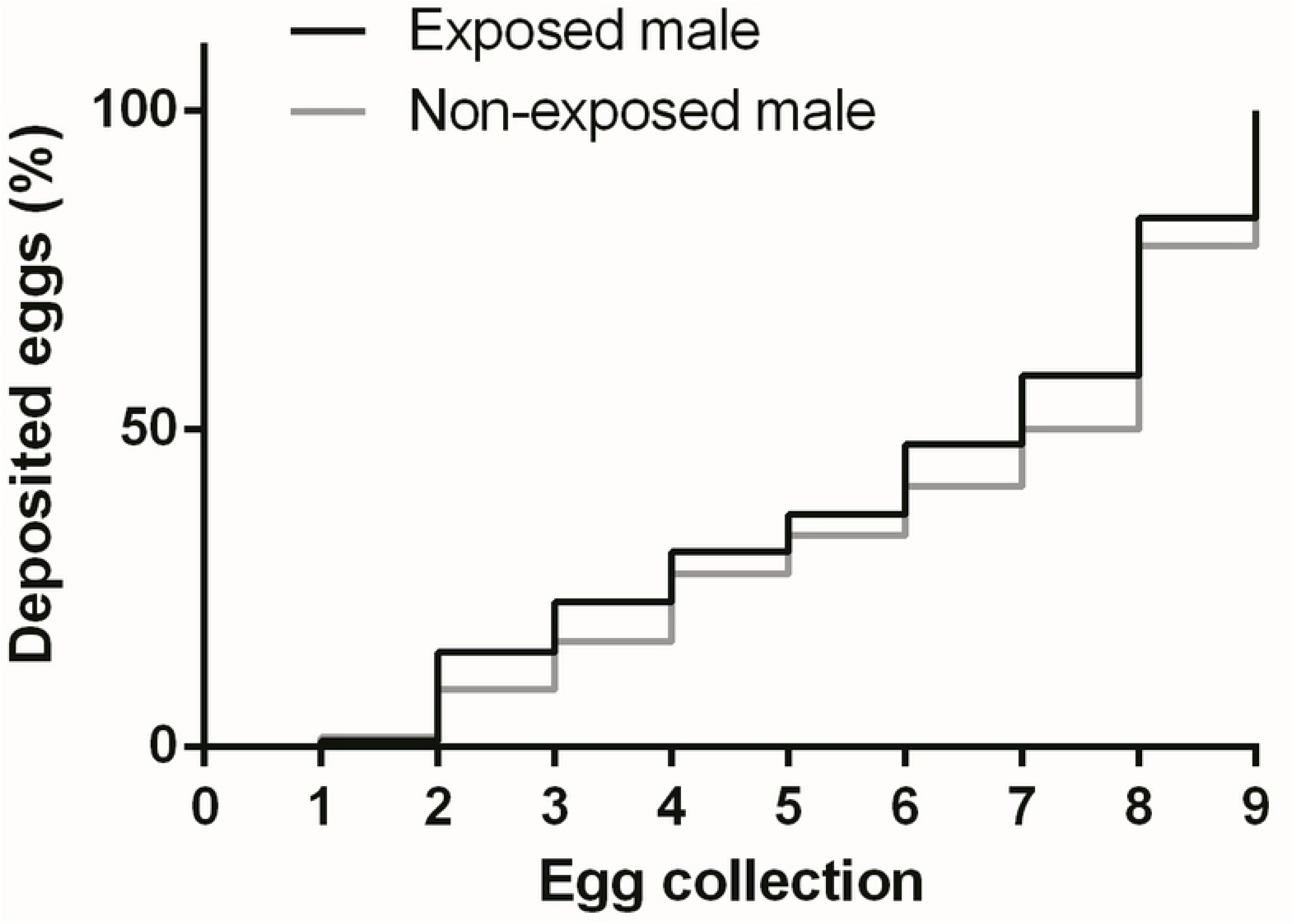
Temporal patterns of egg-laying by females mated to males exposed or non-exposed to guava volatiles.

### Experiment 2. Mating and reproductive output of females offered males treated or not treated with methoprene

Successful matings were recorded in 152 out of 166 mating arenas. Females mated significantly more often with males treated with methoprene than with non-treated males (G = 11.76, N = 152, d.f. = 1, p < 0.001) (Fig 4). However, male type had no effect on any of the measured variables (latency to mate: t = 0.357, d.f. = 150, p = 0.722; mating duration: t = 0.257, d.f. = 146, p = 0.797; fecundity: t = 0.715, d.f. = 38, p = 0.476; fertility: t = 1.100, d.f. = 38, p = 0.278) (Fig 5a-d). Likewise, the temporal pattern of oviposition was independent of the type of male selected by the female (50% of the total eggs deposited by the 5^th^ egg collection for both groups) (χ^2^ = 1.61, d.f. = 1, p = 0.205) (Fig 6).

**Fig 4.**
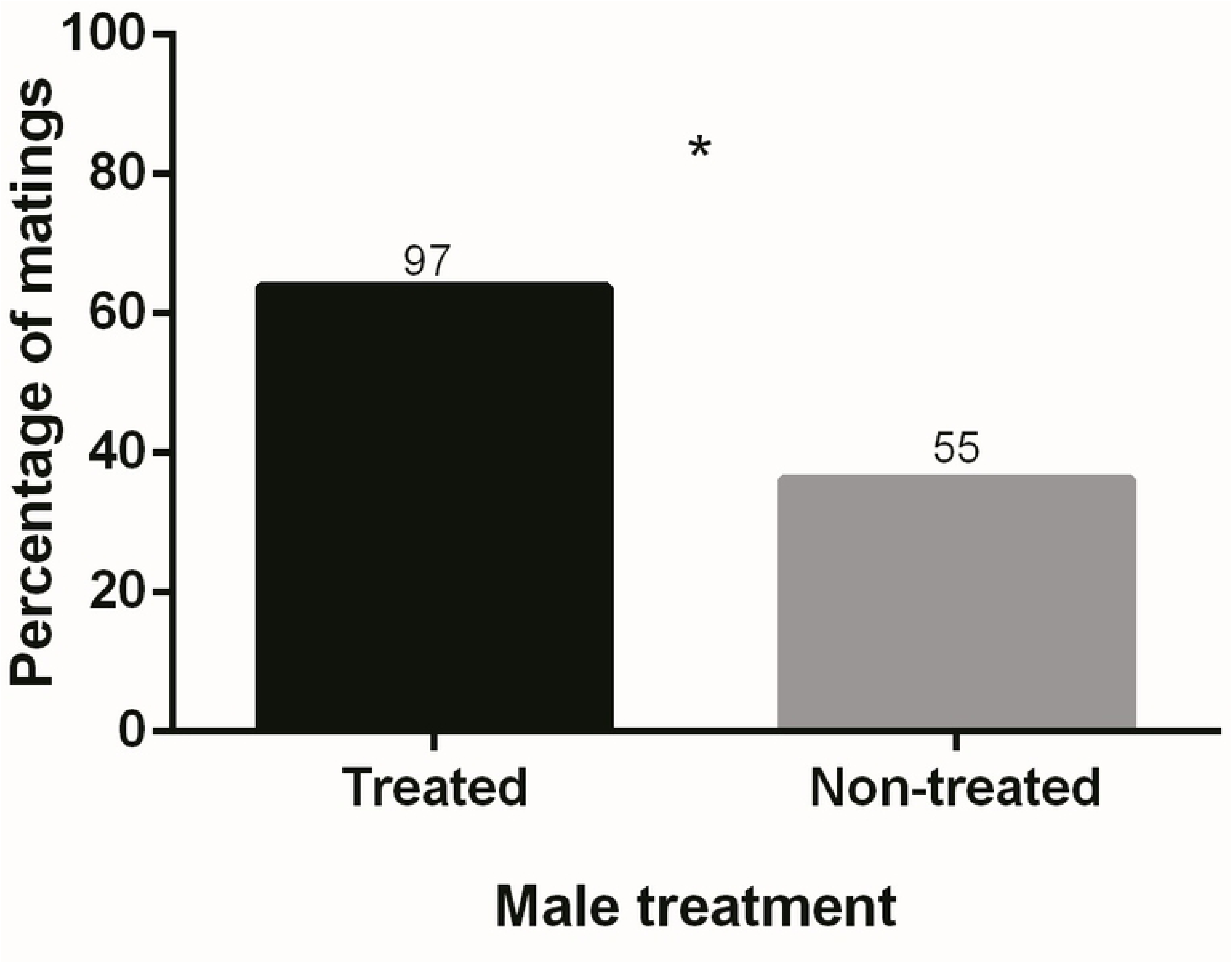
Percentage of matings obtained by males treated or non-treated with methoprene. Numbers above bars represent numbers of matings. * statistically significant difference (α = 0.05).

**Fig 5.**
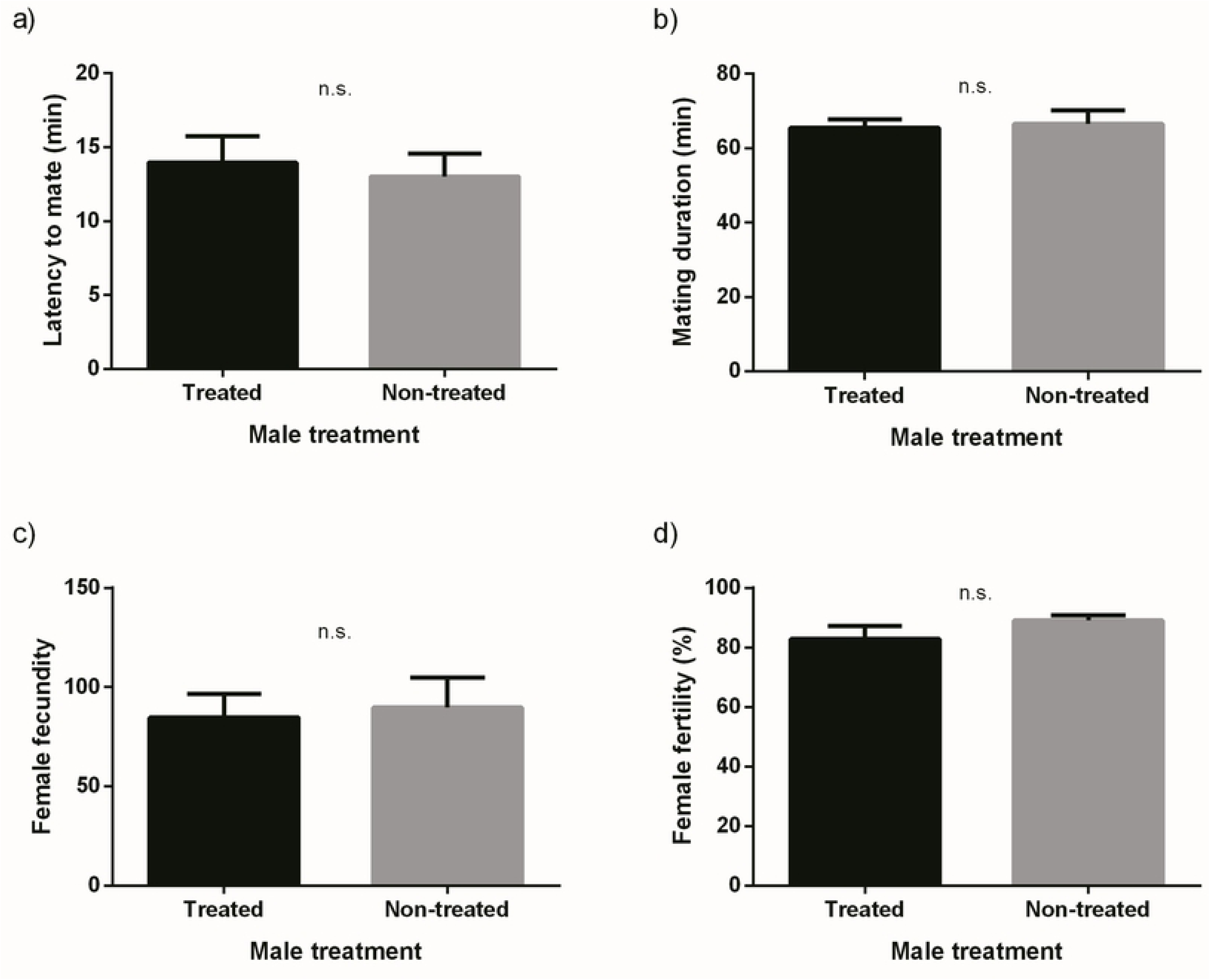
Latency to mate (a), mating duration (b), fecundity (*per capita*) (c) and fertility (d) for females mated to males treated or non-treated with methoprene (mean and SE). n.s. not statistically significant difference (α = 0.05).

**Fig 6.**
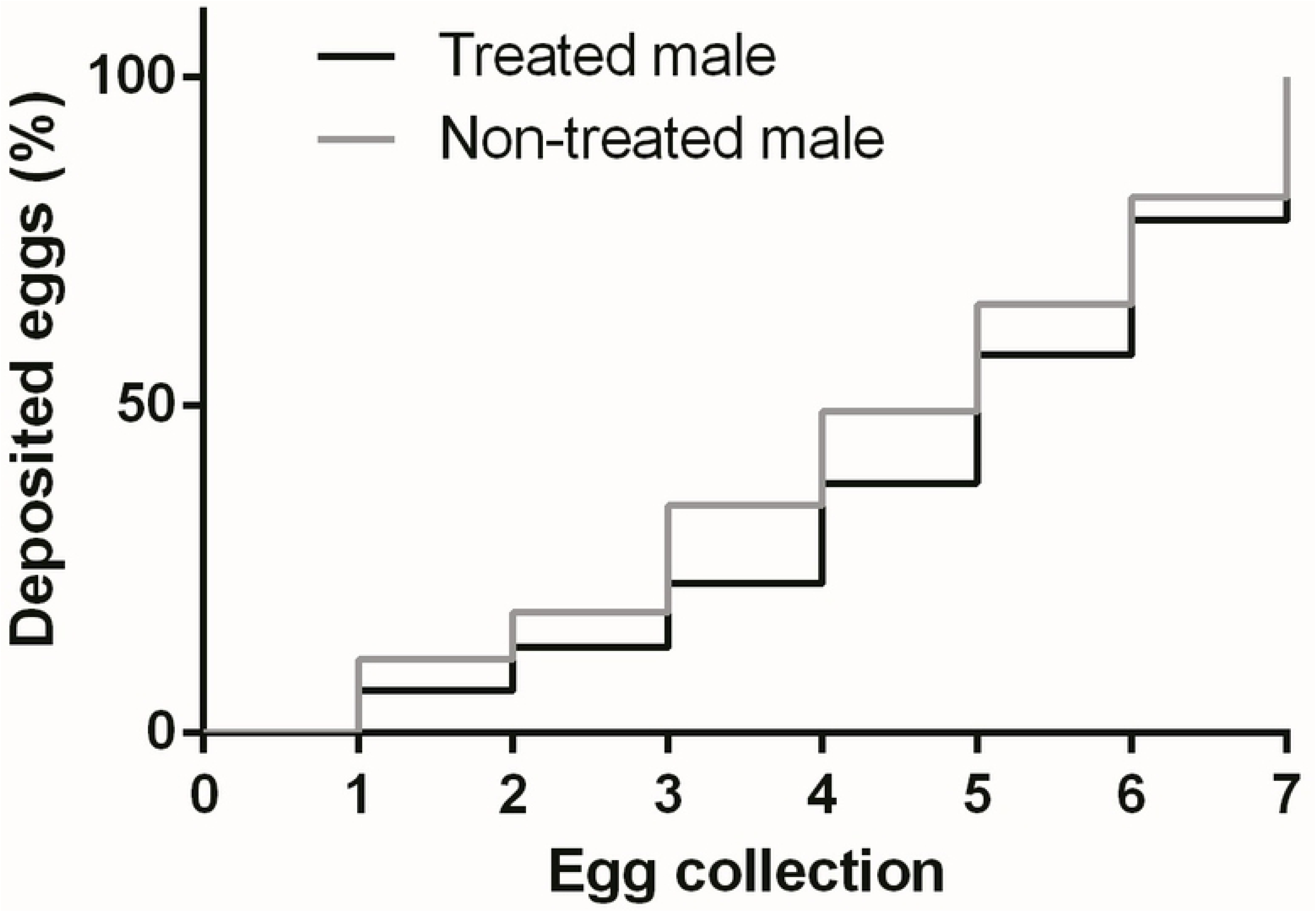
Temporal pattern of egg deposition by females mated to males treated or non-treated with methoprene.

## Discussion

Documenting female preference based on particular male traits is more easily, and thus more frequently, accomplished than demonstrating fitness benefits to females accruing from such preferences. Vera *et al*. [42] and Bachmann *et al*. [43,44] observed that *A. fraterculus* females prefer males that had been exposed to volatiles of guava fruit (one of its main hosts) or treated with methoprene (an artificial analogue of JH). Here, we verified these patterns of sexual selection and investigated whether female preferences can be explained on the basis of a direct fitness benefit. We found that females that selected males exposed to guava volatiles had higher fecundity than those selecting non-exposed males. Therefore, such enhanced egg output is consistent with the existence of a direct fitness benefit. In other words, preference for males exposed to guava volatiles represents an adaptive, fitness-based decision. Additionally, matings that involved guava exposed males lasted longer, which, as discussed below, may also be considered beneficial for the females. On the other hand, for methoprene treatment, we did not detect any association between preference and fitness benefits for females. Female latency to mate did not differ with male status for both guava volatile and methoprene treatments. Thus, although exposure to guava volatiles and application of methoprene boosted male mating success, neither treatment stimulated more rapid mating decisions by females.

The difference in fecundity between females mated to guava-exposed and non-exposed males was constant over time, which indicates that the benefit derived from mating with exposed males was long-lasting and not restricted to the beginning of the oviposition period. The higher fecundity observed here is consistent with Kumaran *et al*. [38], who recorded increased fecundity in *B. tryoni* females mated to males that were previously exposed to zingerone, a floral compound produced by orchids, that enhaces male mating success. Particularly, our findings constitute the first evidence of such mechanism involving volatiles from the host fruit. For tephritids, that study and the present one provide the only clear evidence of a direct benefit associated with female mate choice, where female preference is mediated by male exposure to plant-derived compounds. Other related studies on tephritids found no support for this phenomenon [22]. For example, females of *B. dorsalis* mate preferentially with methyl eugenol-fed males, but they did not show an increase in their fecundity, fertility, or survival [37,65]. Similarly, *C. capitata* females did not exhibit any reproductive or survival benefits after mating with preferred, ginger root oil treated males [13].

One mechanism by which fecundity might have increased is the acquisition of higher quality male accessory gland products (AGPs) during copulation, which occurs in many insects including tephritids [66–68]. There is a large variety of physiologically active substances, such as proteins and juvenile hormone in the ejaculate, that may inhibit female remating propensity and induce egg maturation [66,69,70]. The perception of guava volatiles by males could act as an indicator of host availability and alter male reproductive physiology, stimulating the production of AGPs that would be transferred in higher amounts to the females with their concomitant impact on fecundity. The possibility that males spend more energetic sources in reproduction (e.g. signalling and AGPs production) when a host is present can be a reasonable explanation, but it is, of course, conjectural, and studies on the effects of guava exposure on male reproductive physiology are required.

In contrast to the guava treatment, mating with methoprene-treated males did not result in increased fecundity, fertility, or copulation duration. The lack of evidence of direct benefits can be interpreted in different ways. First, because methoprene is a synthetic compound, it may be possible that looking at the behavior of females from an evolutionary perspective (i.e. relating their preference with direct fitness benefits) is misleading. In this scenario, methoprene would induce a higher rate of pheromone release in males, which triggers female acceptance over non-treated males [44] by exploiting females’ sensory channel with no reward in terms of their own potential fecundity. However, juvenile hormone analogues do exist in nature [45–47] and some insects consume them, increasing their reproductive output [48]. Kumaran *et al*. [38] found that females of *B. tryoni* obtained a direct benefit by mating males exposed to zingerone (a natural compound) as well as males exposed to cuelure (a synthetic compound). This argues against the idea that the lack of benefits associated to methoprene lays on the fact that methoprene is a synthetic compound. Methoprene is similar to JH in chemical structure and, more importantly, its role on *A. fraterculus* seems to replicate that of JH [55,71,72]. Therefore, the strong preference of females for methoprene-treated males could, still, be associated to other benefits, yet unidentified. First, because males treated with methoprene release more pheromone than no-treated males, females could be obtaining a direct benefit if treated males are more easily located, consequently reducing predation risks and mate location costs [13]. Second, indirect (genetic) benefits may be involved in female preference for methoprene-treated males. In *B. tryoni*, Kumaran *et al*. [39] found that the sons of males that were treated with cuelure and zingerone detected and located these chemicals more effectively than sons of non-exposed males, thus showing evidence of a potential “sexy son” mechanism. Potential indirect benefits, like those reported for *B. tryoni* [38,39], should also be studied for *A. fraterculus* females mated not only with methoprene-treated males but also in females mated with guava-exposed males, as direct and indirect effects can both be influencing female choice.

Copula lasted longer for guava exposed males than for non-exposed ones. *Anastrepha fraterculus* females appear to remate primarily to replenish sperm [73], and this tendency is negatively associated to copula duration [49]. Thus, long copulations may be adaptive, because the transfer of large amounts of sperm would eliminate or delay the expenditure of energy and time to find males. Because predation risk increases during courtship and mating [50] delaying remating would also be beneficial in terms of survival. These benefits may, of course, be offset to some degree by any increase in predation risk during a single but lengthy copulation and the fact that overall genetic variability of the progeny is expected to be lower. In the case of methoprene, copula duration did not differ between treated and non-treated males. This agrees with Haq et al. [19] for *Z. cucurbitae* and shows a general lack of other types of benefits associated to methoprene.

The preference of females for males treated with sexual enhancers, together with a resulting increased fecundity could have practical implications in the context of SIT. In a mass rearing facility, an increase in fecundity would mean a higher yield which directly translates into lower costs of maintenance (because fewer reproductive adults would be needed). Also, if enhanced, but sterile, males were able to induce sterility in wild females, then a higher proportion of the reproductive output of the wild population would be unviable. Nonetheless, in order to extend our results to the context of SIT, further research is needed. For instance, the impact of irradiation on both the enhancement of males mating success and females fecundity were not assessed.

To conclude, previous work [43] showed that the preference of females for males exposed to guava volatiles could be explained at a proximal level by a higher rate of sexual displays and sex pheromone release by exposed males. Here, at least for guava volatiles, we found evidence that female preference could also be explained at an evolutionary level, because females that mate with guava-exposed males obtained a direct benefit in terms of increased fecundity. Our results contribute to a better understanding of the mechanisms related to mate choice and their evolutionary implications. However, the ultimate (physiological) causes of an increased fecundity after mating with a sexually enhanced male still remain unknown and represent an interesting field of study. It is worth mentioning that, in all the experiments of the present study, females were fed with diet of high nutritional quality and presumably met, to a large extent, their nutritional and physiological needs (e.g., oocytes development). Aluja et al. [11] found that *Anastrepha ludens* and *Anastrepha obliqua* females fed on protein diets developed more oocytes than those fed without protein. It is thus possible that a rich diet actually reduced the positive effects of mating with guava-exposed males and that females feeding on a lower quality diet, as expected in the wild, would show an even greater increase in fecundity than found here. The interaction between female nutritional status and potential benefits gained through mate choice should be considered in future studies.

## Acknowledgements

The authors are grateful to our colleagues at IGEAF, INTA, for their collaboration during the tests. These studies were funded by the FAO/IAEA and Ministerio de Ciencia, Tecnología e Innovación Productiva of Argentina (through Research Contract 16483 and FONCYT PICT 2013 - 054 to JLC and DFS, respectively).

